# Standardizing microsatellite panels for individual identification of seabirds’ Snow Petrel *Pagodroma nivea* and Wilson’s Storm Petrel *Oceanites oceanicus* in Antarctica

**DOI:** 10.1101/221861

**Authors:** Anant Pande, Nidhi Rawat, Kuppusamy Sivakumar, Sambandan Sathyakumar, Vinod B. Mathur, Samrat Mondol

## Abstract

Seabirds are known to be important indicators of marine ecosystems health. Procellariiformes are one of the most abundant seabird species distributed from warm tropical to cold temperate regions including Antarctica. With few long-term studies on breeding seabirds at the Antarctic continent, crucial biological parameters such as genetic variation, population genetic structure and past population demography is lacking for most of the commonly occurring species. Under the ‘Biology and Environmental Sciences’ component of the Indian Antarctic programme, long-term monitoring of Antarctic biodiversity is being conducted. In this paper, we describe a panel of 12 and 10 cross-species microsatellite markers for two relatively less studied seabird species in Antarctica, snow petrel *Pagodroma nivea* and Wilson’s storm petrel *Oceanites oceanicus*, respectively. These loci showed high amplification success and moderate level of polymorphism in snow petrel (mean no. of alleles 7.08±3.01 and mean observed heterozygosity 0.35±0.23), but low polymorphism in Wilson’s storm petrel (mean no. of alleles 3.9±1.3 and mean observed heterozygosity 0.28±0.18). The results demonstrate that these panels can unambiguously identify individuals of both species from various types of biological materials. This work forms a baseline for undertaking long-term genetic research of Antarctic seabird species and provides critical insights into their population genetics.

## INTRODUCTION

Seabirds, being top predators, maintain structure of marine food webs, regulate island and marine ecosystem processes and act as indicators of marine ecosystem health (Lascelles *et al*. 2012; Paleczny *et al*. 2015). Their natural ability to fly over large distances, extreme life history strategies (monogamy, slow reproduction, late sexual maturity), natal philopatry, high visibility and dependence on land for breeding makes them ideal candidates for long-term population level studies (Piatt *et al*. 2007). A number of recent studies focusing on seabird population monitoring have highlighted the threatened status of seabirds across the globe (Croxall *et al*. 2012), especially in the southern ocean where seabird populations have declined substantially over last few decades (Paleczny *et al*. 2015). This has led to efforts focusing on understanding seabird population dynamics using interdisciplinary approaches to aid conservation and management across their distribution range (Croxall *et al*. 2012; Taylor and Friesen 2012).

Among seabirds, order Procellariiformes includes Petrels, Shearwaters, Albatrosses, Storm Petrels and Diving Petrels representing the most widely distributed and abundant species (Warham 1996). In spite of their wide range and large population sizes, long-term ecological and genetic data exists for few of these species across the globe. In addition to several ecological studies on Procellariiformes (Croxall *et al*. 2012), some recent studies have used genetic data to address important biological parameters such as relatedness, population structure, past population demography (e.g. see Gómez-Díaz *et al*. 2009 for Cory’s shearwater; Welch *et al*. 2012 for Hawaiian petrel) for species distributed in tropical and arctic marine ecosystems. Research on Procellariiformes’ biology is relatively limited in the Southern Ocean ecosystem, especially in Antarctica because of its remoteness and associated logistical difficulties. Despite few site-specific monitoring of some Procellariiformes on sub-Antarctic islands (e.g. Brown *et al*. 2015 for giant petrels; Quillfeldt *et al*. 2017 for Antarctic prion, thin-billed prion and blue petrel) and Antarctic coast (e.g. Barbraud & Weimerskirch, 2001 on snow petrel; Barbraud and Weimerskirch 2006 on multiple species; Techow *et al*. 2010 on giant petrels), long-term ecological as well as genetic research is sparse. Nunn and Stanley (1998) reported the phylogenetic relationships among procellariform species’ using a neighbour-joining approach, but within each family groups detailed population genetic information is lacking. Other preliminary studies have used Restriction Fragment Length Polymorphisms (RFLP) and allozymes to investigate genetic variation and extra-pair paternity in snow petrel as well as some other Procellariiformes (Jouventin and Viot 1985, Viot *et al*. 1993, Lorensten *et al*. 2000, Quillfeldt *et al*. 2001) in Antarctica.

The ‘Biology and Environmental Sciences Programme’ of Indian Scientific Expeditions to Antarctica has a specific focus on understanding distribution, abundance, population dynamics and genetics of Antarctic seabirds, including Procellariiformes. As part of this program, comprehensive ecological surveys were conducted between 2009-2016 to understand seabird and marine mammal ecology around Indian Antarctic research stations (Pande *et al*. 2017). Currently this programme is focused on generating baseline genetic data of breeding seabird species found around Indian area of operations in Antarctica, especially on Snow Petrel *Pagodroma nivea* and Wilson’s Storm Petrel *Oceanites oceanicus*. Snow petrel is endemic to Antarctica and Southern Ocean with breeding distribution along Antarctic coast including some inland mountains and few sub-Antarctic islands (Croxall *et al*. 1995). On the other hand, Wilson’s storm petrel has a much wider breeding distribution from Cape Horn to the Kerguelen Islands and coastal Antarctica. It also migrates to the midlatitudes of the north Atlantic, north Indian and Pacific Ocean during non-breeding period (BirdLife International 2017). Effective monitoring of these species in the Indian Antarctic sector will require systematic information on their distribution, current population status and genetics. With the broad objective of assessing population genetic structure, we describe a panel of cross-species microsatellite markers for individual identification of snow petrel and Wilson’s storm petrel in Antarctica and sub-Antarctic islands. These tested panels will be of great help in understanding genetic variation, genetic relatedness and demographic history of both these species across their ranges.

## METHODS

### Permits and ethical clearances

All samples were collected under the ‘Biology and Environmental Sciences’ component (Letter no: NCAOR/ANT/ASPA/2014-15/01) of the Indian Scientific Expeditions to Antarctica with appropriate approvals from the Environment Officer, Committee for Environmental Protection (Antarctic Treaty Secretariat), National Centre for Antarctic and Ocean Research, Earth System Science Organisation, Ministry of Earth Sciences, Government of India, Goa, India.

### Study Area

Sampling was carried out at Larsemann hills, Prydz bay and Schirmacher oasis, Central Dronning Maudland (Figure 1); close to permanent Indian research stations in Antarctica *Bharati* (Larsemann hills) and *Maitri* (Schirmacher oasis). Distance between these two study areas is about 2,500 km. Larsemann hills (69° 20’S to 69° 30’S; 75° 55’E to 76° 30’E coordinates), are a group of islands in Prydz Bay located on the Ingrid Christensen Coast, Princess Elizabeth Land of east Antarctica. It comprises of variously sized islands and peninsulas, located halfway between the eastern extremity of the Amery Ice Shelf and the southern boundary of the Vestfold Hills. Schirmacher Oasis, Central Dronning Maudland (70° 44’ to 70° 46’ S and 11° 22’ to 11° 54’ E coordinates) is situated on the Princess Astrid coast about 120 km from the Fimbul ice shelf. Four species of seabirds (Adelie penguin *Pygoscelis adeliae*, south polar skua *Stercorarius maccormickii*, snow petrel and Wilson’s storm petrel) breed in the ice-free areas of Larsemann hills whereas only south polar skua breeds at Schirmacher oasis (Pande et al. 2017). Other key wildlife species found around these areas include emperor penguin *(Aptenodytes forsteri)*, crabeater seal *Lobodon carcinophaga*, leopard seal *Hydrurga leptonyx*, Ross seal *Ommatophoca rossii*, Weddell seal *Leptonychotes weddellii* and orca *Orcinus orca*.

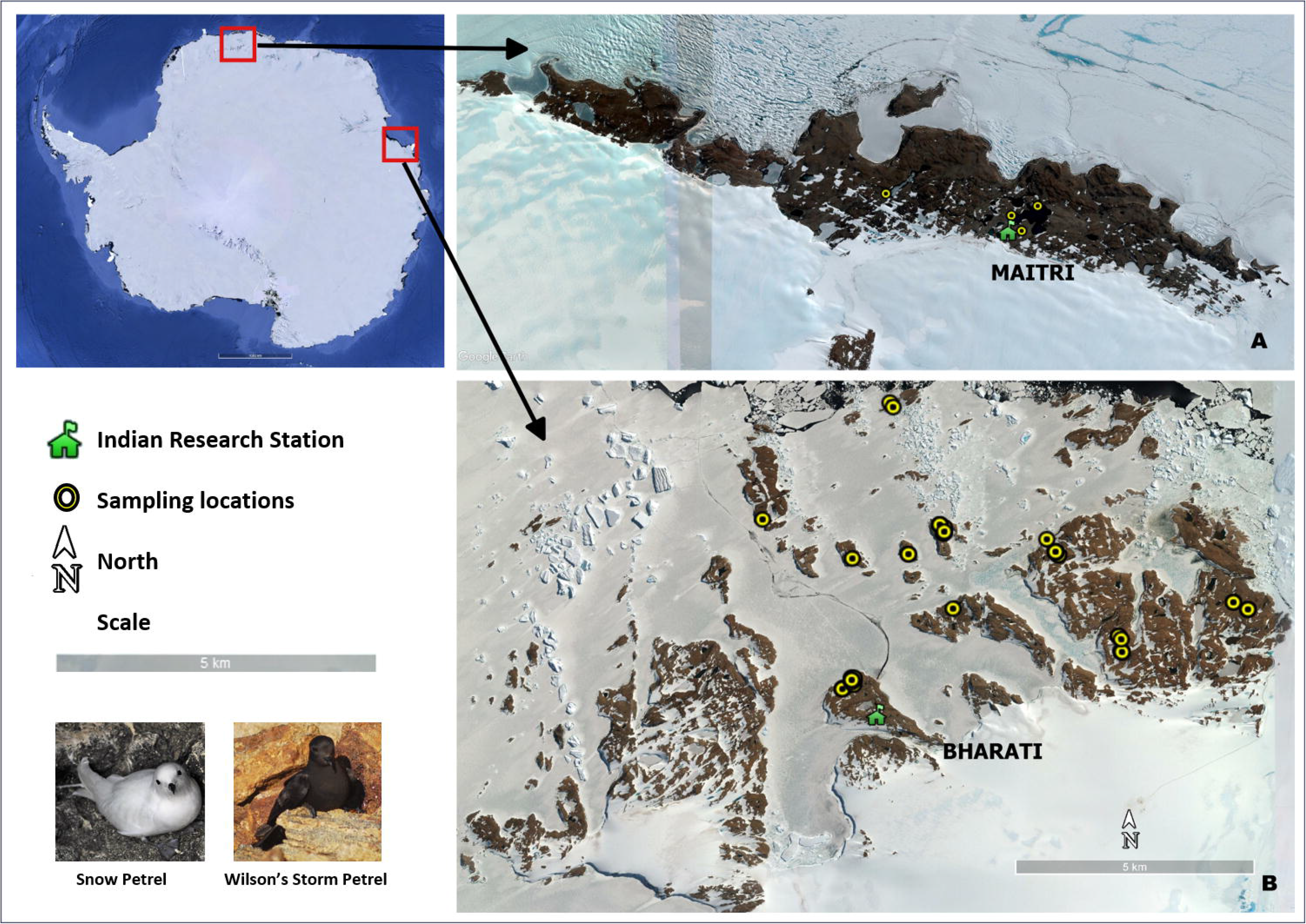

### Field Sampling

Sampling for this study was conducted as part of the ‘Antarctic Wildlife Monitoring Programme’ under the Indian Scientific Expedition to Antarctica (Expedition nos. 33, 34 and 35)’ during the austral summers (November-March) of 2013-14, 2014-15 and 2015-16. We adopted a systematic genetic sampling approach under the seabird nest monitoring protocol (see Pande *et al.*, 2017) for snow petrel sample collection. First, identified nest sites with breeding snow petrel individuals were selected for genetic sampling. Subsequently, both nondestructive (buccal swabs and blood smears) and non-invasive (hatched eggshells and abandoned eggs) sampling approaches were used to collect biological materials from monitored nesting sites. During non-destructive sampling of snow petrel individuals, birds were carefully hand-captured at their nest cavities and buccal swabs or blood samples were collected. Blood samples were collected from bird’s brachial vein using 0.1 ml sterilized syringe needles and stored in an EDTA containing vial. All individuals were released within 60 seconds of capture. We could also collect few hatched eggshells and abandoned eggs from the nests. In addition, opportunistic sampling of snow petrel carcasses was also conducted. These dead animals were mostly predated by south polar skua or naturally dead. Systematic snow petrel sampling was conducted only at Larsemann hills. No nesting sites of snow petrel were found at Schirmacher oasis during field surveys but opportunistic sampling of carcasses was conducted.

Similarly, Wilson’s storm petrel tissue samples were collected from monitored nesting sites at Larsemann hills. All genetic samples of Wilson’s storm petrel were collected opportunistically through carcass tissue collection as capturing them was not possible due to their narrow nest cavities. No Wilson’s storm petrel samples were collected from Schirmacher oasis. Samples collected at field were stored at -20° C at respective Indian Antarctic research stations before being brought to Wildlife Institute of India, Dehradun for further laboratory analysis.

### Primer selection

As there is no published work available with nuclear DNA markers for snow petrel and no species-specific microsatellite markers are yet developed, we tested a panel of cross species markers for individual identification of snow petrels. We selected a total of 15 microsatellite markers earlier developed for Hawaiian petrel *Pterodroma sandwichensis* (Nine markers from Welch and Fleischer 2011) and white-chinned petrel *Procellaria aequinoctialis* (Six markers from Techow and O’Ryan 2004). These markers were selected based on their polymorhic information content (number of alleles as well as expected heterozygosity) in the aforementioned species.

On the other hand, a set of cross-specific microsatellite markers developed for prion species has been tested in Wilson’s storm petrel (Moodley *et al*. 2015). However, the study has reported very low amplification success rate. In this study, we also tested these 15 microsatellite loci for individual identification of Wilson’s storm petrel.

### DNA extraction and primer standardization

We used tissue samples of snow petrel and Wilson’s storm petrel for initial standardization and validation of microsatellite panel. Genomic DNA was extracted in duplicate from all tissue samples using commercially available DNeasy Tissue kit (QIAGEN Inc.) using a modified approach. In brief, all samples were macerated with sterile blades independently, followed by overnight complete tissue digestion with 25 μl proteinase-K. Post-digestion, extraction was performed using Qiagen animal tissue spin column protocol. DNA was eluted twice with 100 μl of 1X TE and stored in −20 °C until further processing. Each set of 11 extractions was accompanied with one extraction control to monitor possible contamination.

All initial PCR standardizations were conducted using tissue DNA samples. Amplifications were carried out for each primer in 10 μl reaction volumes containing 4 μl Qiagen Multiplex PCR buffer mix (Qiagen Inc.), 0.2 μM labeled forward primer, 0.2 μM reverse primer, 4 μM BSA and 2 μl of 1:10 diluted DNA extract. The temperature regime included an initial denaturation (94 ° C for 15 min); 35 cycles of denaturation (94 ° C for 30 s), annealing (53 or 57 °C for 45 s) and extension (72 °C for 45 s); followed by a final extension (72 °C for 30 min). Post-temperature standardization, primers with identical annealing temperatures was optimized for multiplex reactions with the same samples of both species (see Table 1). Subsequently, all test samples were amplified with standardized parameters. During all amplifications, both extraction controls and PCR negative controls (one PCR negative every set of amplifications) were included to monitor any possible contamination. PCR products were visualized in 2% agarose gels, and genotyped using LIZ500 size standard in an automated ABI3500XL genetic analyzer. Microsatellite alleles were scored using program GENEMARKER (SOFTGENETICS Inc.) and allele bins for each locus were created from the data generated. We randomly re-genotyped 15% of each locus from different samples to check for reliable genotypes and estimated genotyping error rates.

**Table 1.**

| Species | Locus | Nucleotide Repeat nature | Dye | Product size range (bp) | Ta (°C) | Number of alleles | Ho | He | Allelic Range | PID (unrelated) Cumulative | PID (sibs) cumulative | Amplification success (%) | Allelic dropout (%) | Multiplex sets for PCR |
| --- | --- | --- | --- | --- | --- | --- | --- | --- | --- | --- | --- | --- | --- | --- |
| Snow Petrel (n=55) | Ptero01 | Di | PET | 82-104 | 53 | 5 | 0.33 | 0.32 | 24 | $4.752 \times 10^{-01}$ | $7.072 \times 10^{-01}$ | 98.2 | 0 | Set 1 |
| | Ptero07 | Tetra | FAM | 177-217 | 53 | 8 | 0.53 | 0.66 | 48 | $7.599 \times 10^{-02}$ | $3.266 \times 10^{-01}$ | 98.2 | 3.6 | |
| | Paequ03 | Di | VIC | 219-243 | 53 | 12 | 0.68 | 0.72 | 24 | $8.205 \times 10^{-03}$ | $1.362 \times 10^{-01}$ | 98.2 | 0 | |
| | Ptero03 | Di | FAM | 165-177 | 53 | 4 | 0.10 | 0.23 | 22 | $4.917 \times 10^{-03}$ | $1.067 \times 10^{-01}$ | 100 | 0 | |
| | Paequ08 | Di | PET | 215-223 | 53 | 4 | 0.16 | 0.18 | 8 | $3.291 \times 10^{-03}$ | $8.799 \times 10^{-02}$ | 100 | 0 | |
| | Paequ02 | Di | PET | 180-200 | 53 | 7 | 0.03 | 0.30 | 30 | $1.642 \times 10^{-03}$ | $6.379 \times 10^{-02}$ | 98.2 | 1.8 | Set 2 |
| | Ptero04 | Di | FAM | 117-147 | 53 | 11 | 0.67 | 0.63 | 32 | $2.906 \times 10^{-04}$ | $3.059 \times 10^{-02}$ | 100 | 0 | |
| | Ptero08 | Tetra | VIC | 181-221 | 53 | 11 | 0.49 | 0.73 | 52 | $2.742 \times 10^{-05}$ | $1.253 \times 10^{-02}$ | 96.4 | 0 | |
| | Paequ10 | Di | NED | 159-183 | 53 | 7 | 0.20 | 0.56 | 12 | $6.217 \times 10^{-06}$ | $6.593 \times 10^{-03}$ | 98.2 | 0 | |
| | Paequ13 | Di | PET | 144-150 | 57 | 4 | 0.07 | 0.44 | 6 | $2.189 \times 10^{-06}$ | $4.063 \times 10^{-03}$ | 100 | 0 | Set 3 |
| | Paequ07 | Di | FAM | 314-320 | 57 | 3 | 0.30 | 0.40 | 6 | $8.665 \times 10^{-07}$ | $2.625 \times 10^{-03}$ | 100 | 0 | |
| | Ptero09 | Tetra | FAM | 161-189 | 57 | 9 | 0.67 | 0.72 | 28 | $1.041 \times 10^{-07}$ | $1.106 \times 10^{-03}$ | 100 | 0 | |
|  | Mean (SD) |  |  |  |  | 7.08 (3.01) | 0.35(0.23) | 0.49(0.19) | 24.5(14.5) |  |  |  |  |  |
| Wilson's Storm Petrel (n=24) | Ptero01 | Di | PET | 165-177 | 53 | 4 | 0.17 | 0.44 | 12 | $3.65 \times 10^{-01}$ | $6.19 \times 10^{-01}$ | 100 | 0 | Set 1 |
| | Paequ10 | Di | NED | 181-191 | 53 | 4 | 0.38 | 0.64 | 10 | $7.44 \times 10^{-02}$ | $2.99 \times 10^{-01}$ | 100 | 0 | |
| | Ptero07 | Tetra | FAM | 177-217 | 53 | 6 | 0.42 | 0.76 | 40 | $7.29 \times 10^{-03}$ | $1.18 \times 10^{-01}$ | 100 | 0 | |
| | Paequ03 | Di | VIC | 219-235 | 53 | 5 | 0.21 | 0.39 | 16 | $2.86 \times 10^{-03}$ | $7.73 \times 10^{-02}$ | 100 | 0 | |
| | Ptero03 | Di | FAM | 88-104 | 57 | 2 | 0.17 | 0.35 | 16 | $1.38 \times 10^{-03}$ | $5.37 \times 10^{-02}$ | 91.7 | 0 | Set 2 |
| | Ptero04 | Di | FAM | 127-139 | 57 | 4 | 0.38 | 0.52 | 12 | $4.30 \times 10^{-04}$ | $3.05 \times 10^{-02}$ | 100 | 0 | |
| | Paequ13 | Di | PET | 146-148 | 57 | 2 | 0.08 | 0.5 | 2 | $1.62 \times 10^{-04}$ | $1.82 \times 10^{-02}$ | 100 | 8.3 | |
| | Paequ08 | Di | PET | 219-227 | 51 | 3 | 0.21 | 0.47 | 8 | $6.04 \times 10^{-05}$ | $1.10 \times 10^{-02}$ | 100 | 0 | Set 3 |
| | Ptero09 | Tetra | FAM | 173-185 | 61 | 6 | 0.67 | 0.55 | 16 | $1.43 \times 10^{-05}$ | $5.88 \times 10^{-03}$ | 91.7 | 0 | |
| | Paequ07 | Di | FAM | 312-318 | 51 | 3 | 0.08 | 0.16 | 6 | $1.03 \times 10^{-05}$ | $5.01 \times 10^{-03}$ | 100 | 4.2 | |
|  | Mean (SD) |  |  |  |  | 3.9 (1.3) | 0.28 (0.18) | 0.48 (0.15) | 13.8(9.7) |  |  |  |  |  |

### Data analysis

Average amplification success was calculated as the percent positive PCR for each locus, as described by Broquet *et al*. (2007). Allelic dropout and false allele rates were quantified manually as the number of dropouts or false alleles over the total number of amplifications, respectively (Broquet *et al*. 2007). We also calculated the Probability of Identity for siblings (PID)sibs, the probability of two individuals drawn from a population sharing the same genotype at multiple loci (Waits *et. al* 2001) using program GIMLET (Valière 2002). We tested the frequency of null alleles in our data set using FREENA (Chapuis & Estoup 2007) whereas summary statistics and tests for deviations from Hardy-Weinberg equilibrium were calculated for each locus using program ARLEQUIN v.3.1 (Excoffier and Schneider 2005).

### Results and discussion

We genotyped a total of 55 snow petrel and 24 Wilson’s storm petrel samples to test and standardize the selected microsatellite markers. Snow petrel samples were selected from blood (n=1), buccal swab (n=2), carcass tissue (n=24) and eggshells (n=28) to test amplification success from different types of biological samples. Wilson’s storm petrel samples were all from muscle tissue of individual carcasses collected in the field.

Of the 15 loci tested during the initial standardization, 12 loci showed amplification for snow petrel (loci Ptero2, Ptero6 and Ptero10 did not amplify), whereas only 10 loci successfully amplified for Wilson’s storm petrel (loci Paequ2, Ptero2, Ptero6, Ptero8 and Ptero10 did not amplify) (see Table 1 for details). Subsequently, these panels of 12 and 10 loci were tested with all snow petrel and Wilson’s storm petrel samples, respectively. For snow petrel, the loci varied from highly polymorphic (Paequ03-12 alleles, H_o_−0.68) to less polymorphic (Paequ13-4 alleles, H_o_−0.07), whereas for Wilson’s storm petrel the loci were moderately polymorphic (Ptero07-6 alleles, H_o_−0.76) to less polymorphic (Paequ13-2 alleles, H_o_−0.08) (see Table 1 for detailed summary statistics). We could not find any locus that deviated from the Hardy-Weinberg Equilibrium, and there was no evidence for linkage disequilibrium between any pair of loci. Overall, the amplification success ranged between 96.4 – 100% for snow petrel and 91.7 – 100% for Wilson’s storm petrel; and allelic dropout rates were 0 – 3.6% and 0 – 8.3% for snow petrel and Wilson’s storm petrel respectively. The estimated cumulative probability of identity assuming all individuals were siblings (PID(sibs)) was found to be 1.1 × 10^−3^ for snow petrel and 5.0 × 10^−3^ for Wilson’s storm petrel. Average values for observed and expected heterozygosity, number of alleles, allelic range sizes are presented in Table 1. The frequency of null alleles across the loci was observed to be low in both the study species (snow petrel −0.11±0.09 and Wilson’s storm petrel −0.15±0.07).

This paper is the first attempt to use nuclear microsatellite markers to individually identify both snow petrel and Wilson’s storm petrel in Antarctica. Based on the results of this study (PID_(sibs)_ value of 1×10^−3^), it can be ascertained that our standardized 12 microsatellite loci panels are sufficient enough to differentiate among related individuals of snow petrel. However, in case of Wilson’s storm petrel (PID_(sibs)_ value of 5 × 10^−3^) the statistical power is not enough, and additional loci need to be standardized to avoid any possible errors in case of population genetic study of Wilson’s storm petrel. Previously tested microsatellite loci by Moodley *et al*. (2015), though not used for individual identification, could be used along with the current panel of markers to increase the statistical power during individual identification in Wilson’s storm petrel. Overall, our results show that both panels of loci provide unambiguous individuals for respective seabird species in Antarctica.

Molecular genetic analysis has become crucial in understanding levels of genetic differentiation, hybridisation and extinction risk in seabird populations (Taylor and Friesen, 2012). In critical ecosystems such as Antarctica, individual-level genetic data can be a valuable tool to study evolution, adaptation, past events of diversifications and extinctions for wide-ranging seabirds. In this study, we could establish the efficacy of cross-species markers in individual identification of two common Antarctic seabird species. We aim to continue this long-term genetic research under the current Antarctica wildlife monitoring programme by increasing spatio-temporal sampling efforts to understand the population structure, relatedness and other aspects and provide insights to seabird behaviour (monogamy, extrapair paternity etc.) and evolution. This detailed genetic research will also aid long-term ecological monitoring of breeding seabird populations and help informed conservation management of these species in Antarctica.

## Acknowledgements

We thank the National Centre for Antarctic and Ocean Research, Ministry of Earth Sciences for providing all logistic support during the Indian Scientific Expeditions to Antarctica. We are grateful to respective expedition leaders and team member volunteers of 33^rd^, 34^th^ & 35^th^ Indian Scientific Expeditions to Antarctica for their support. We thank A. Madhanraj and MEERCAT lab members for their help in laboratory analysis of samples. We sincerely thank Wildlife Forensics and Conservation Genetics Cell, CAMPA Cell, Research Coordinator and Dean, Wildlife Institute of India for their support. We thank the Wildlife Institute of India and DST-INSPIRE faculty Award to SM (Grant no: IFA12-LSBM-47) for financial support to this study.

